## supplementary Figure 1-12 for "Molecular architecture of the tumor microenvironment caused by *BRCA1* and *BRCA2* somatic mutations in lung adenocarcinoma"

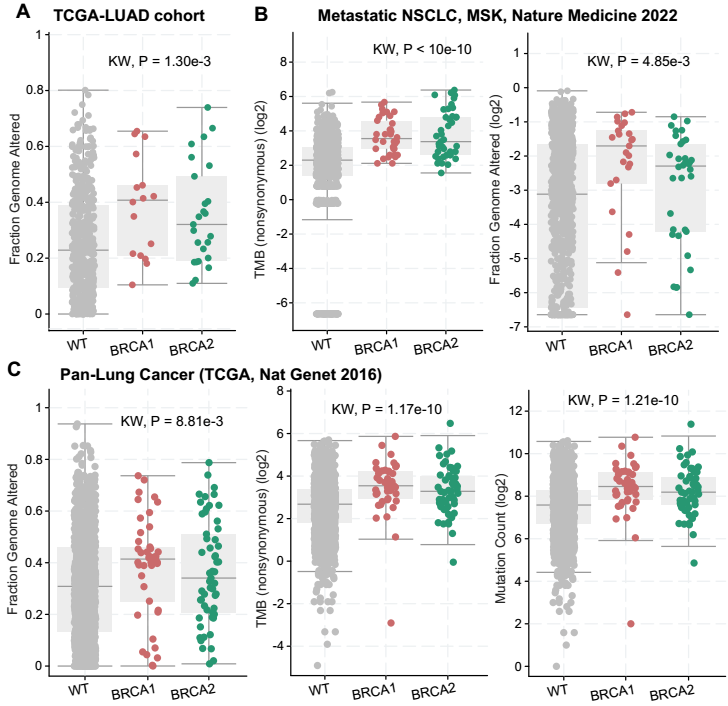

**Fig. S1. *BRCA1* and *BRCA2* mutations are associated with genomic instability. A-C** Distribution of genomic instability indicators (fraction of genome altered, TMB, and mutation count) in LUAD and pan-lung cancer patients with wild-type, *BRCA1* and *BRCA2* mutation.

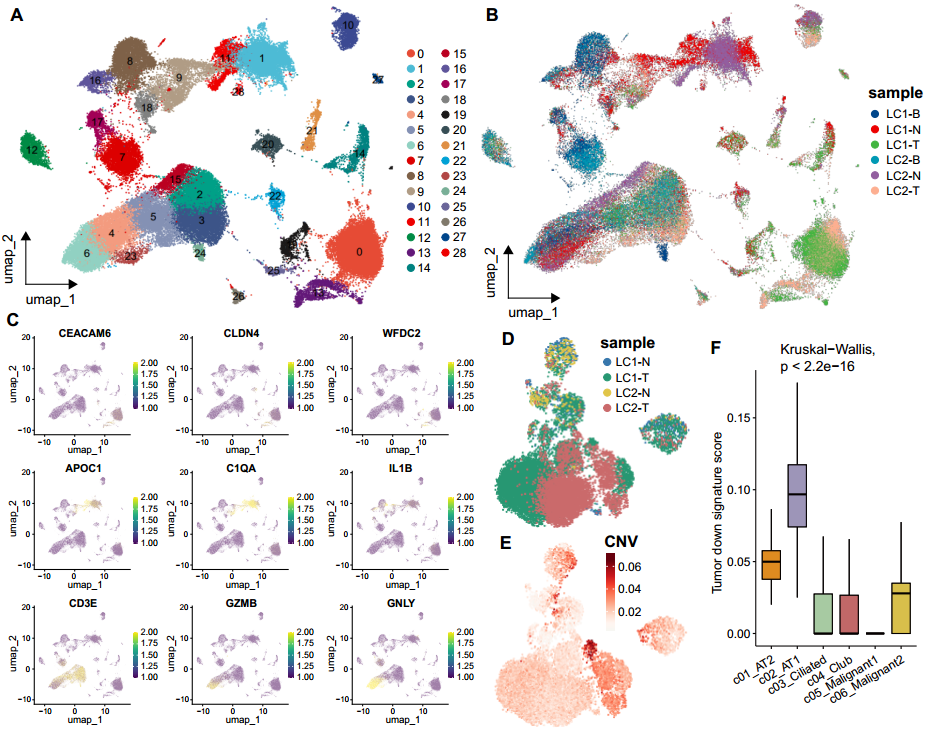

**Fig. S2. Single-cell transcriptome analysis in LUAD patients. A-B** UMAP visualization of cell clusters (**A**) and samples (**B**) in patients with LUAD. **C** The expression of the representative markers. **D-E** UMAP of sample (**D**) and CNV score (**E**) in epithelial and malignant cells. **F** Box plot shows the score of tumor downregulated signature.

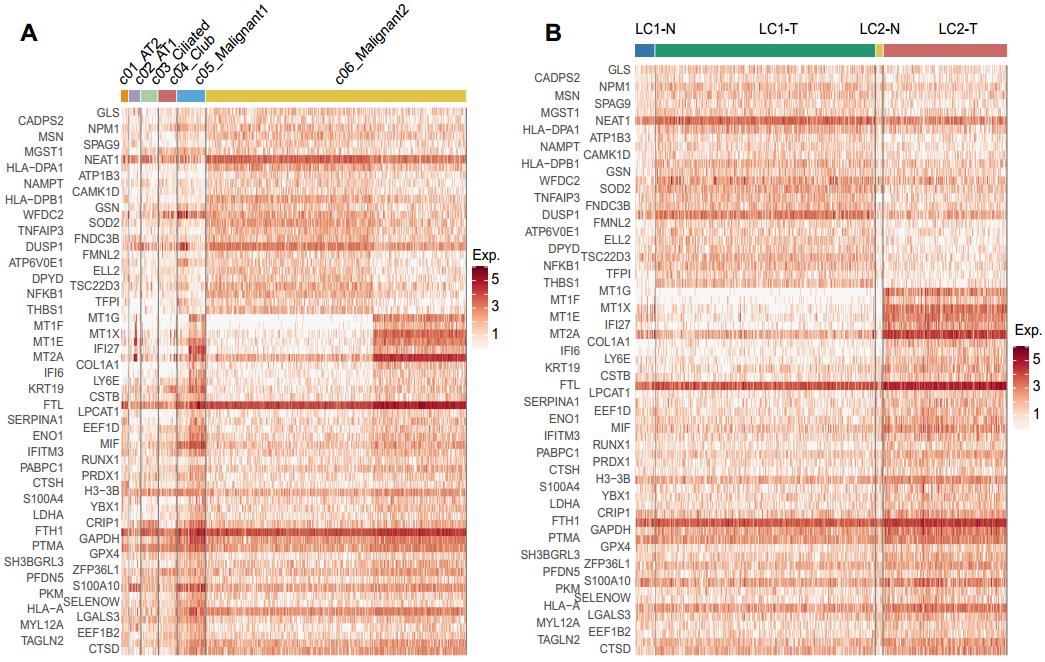

**Fig. S3. The expression of differential genes. A-B** The differential expression genes from malignant cells compared to epithelial cells, was sorted by cell type (**A**) and sample (**B**).

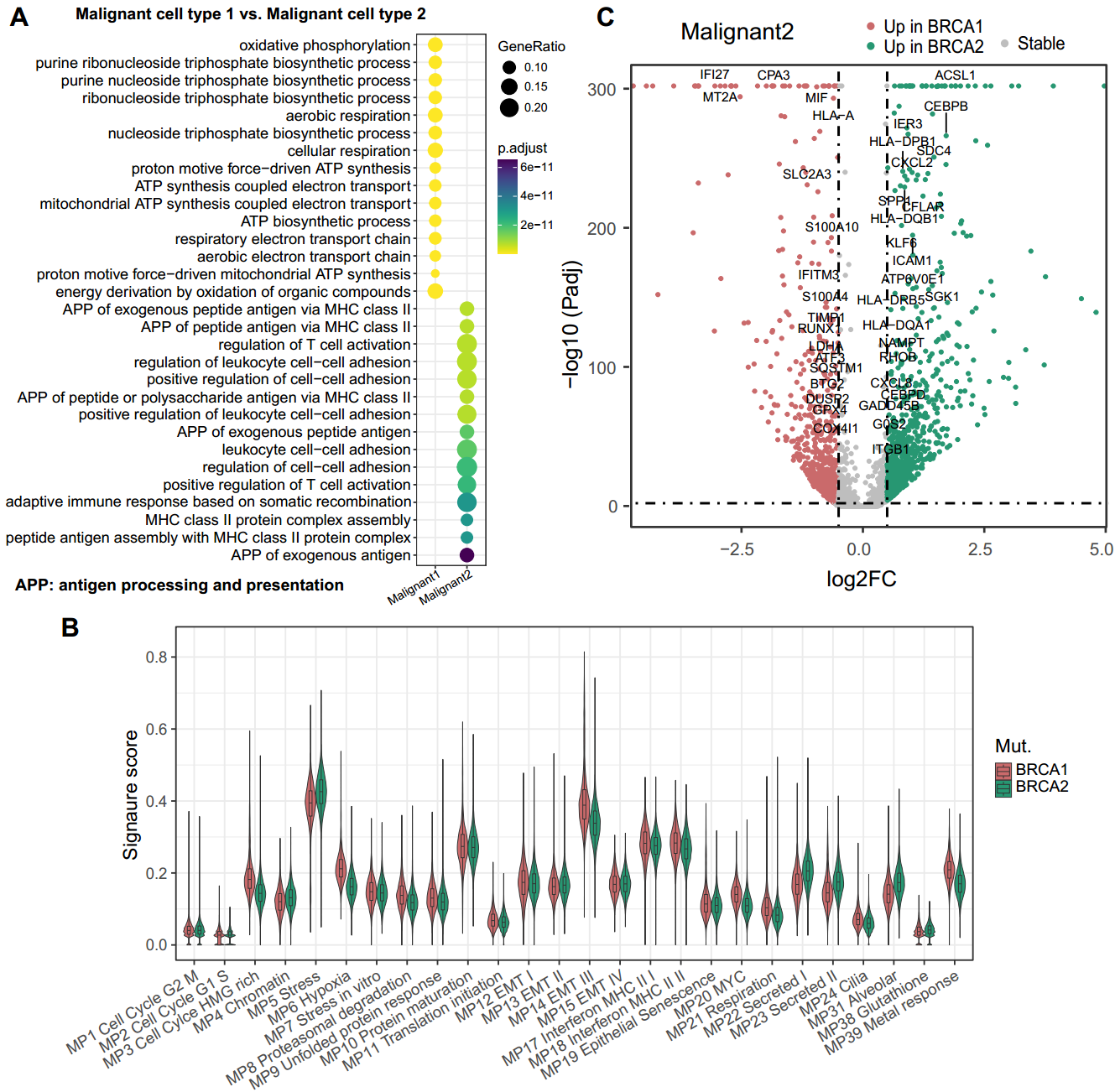

**Fig. S4. Key programs driven by *BRCA1* and *BRCA2* mutations. A** Functional enrichment analysis of differentially expressed genes between two types of malignant subsets. **B** The distribution of known cancer MP score between *BRCA1* and *BRCA2* mutated malignant cells. **C** Volcano diagram of differentially expressed genes between *BRCA1* and *BRCA2* mutated malignant cells (malignant2 cell type).

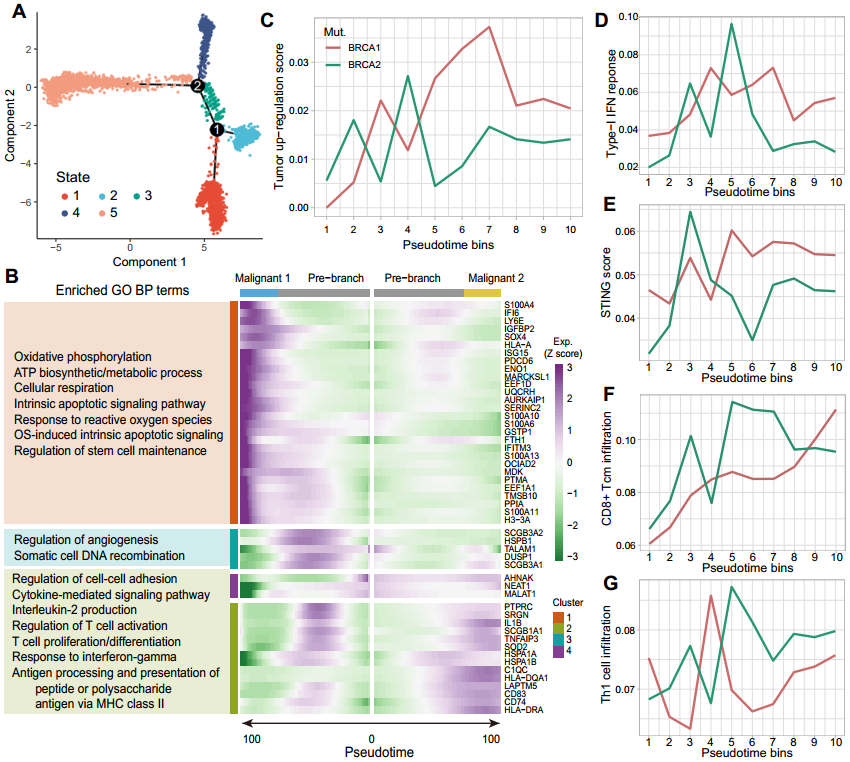

**Fig. S5. Evolutionary analysis of *BRCA1* and *BRCA2* mutant tumors. A** Cell states of trajectory analysis in epithelial and malignant cells. **B** The top 50 cell fate genes and their enriched GO BP terms in two types of malignant cell subsets. **C-G** Pseudotime was broken down into 10 bins to smooth gene expression patterns. Average module score between *BRCA1* and *BRCA2* mutation groups for tumor up-regulation (**C**), type I IFN response (**D**), STING (**E**), and CD8+ Tcm infiltration (**F**) and Th1 infiltration (**G**) gene modules.

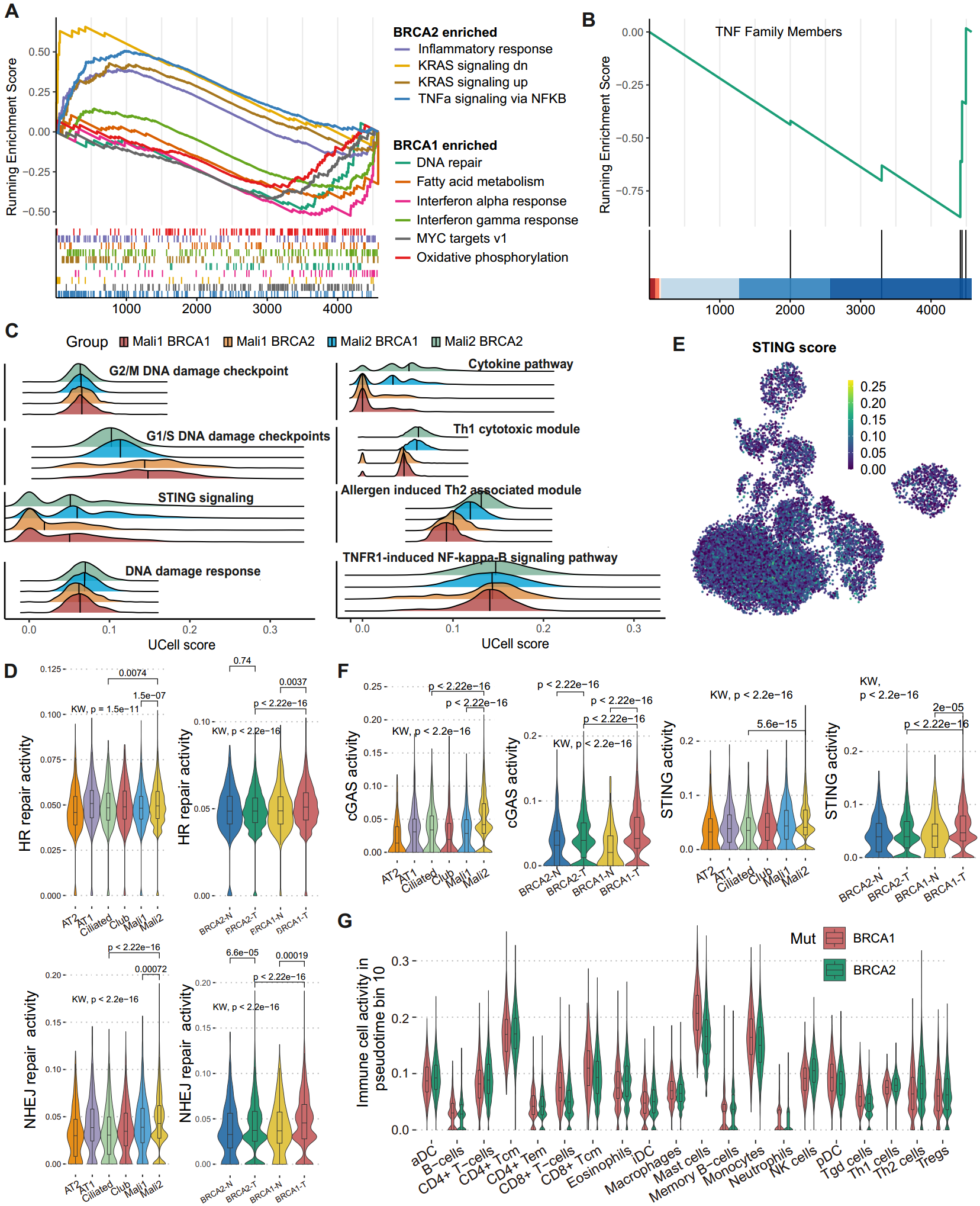

**Fig. S6. Heterogeneity of *BRCA1/2* mutations in tumor lymphoid activity. A-B** GESA plot of hallmarks (**A**) and TNF family members (**B**). NES: normalized enrichment score. Candidate criteria were |NES|>1, P<0.05. **C** Density ridge plot of representative pathways in two types of malignant subset with *BRCA1* and *BRCA2* mutations. **D, F** Activity comparison of and DSBs-related DNA repair pathways (**D**) and cGAS-STING signaling pathways (**F**). **E** UMAP visualization of STING pathway score. **G** Immune cell activities between *BRCA1* and *BRCA2* mutations in bin 10 of malignant cells according to pseudotime.

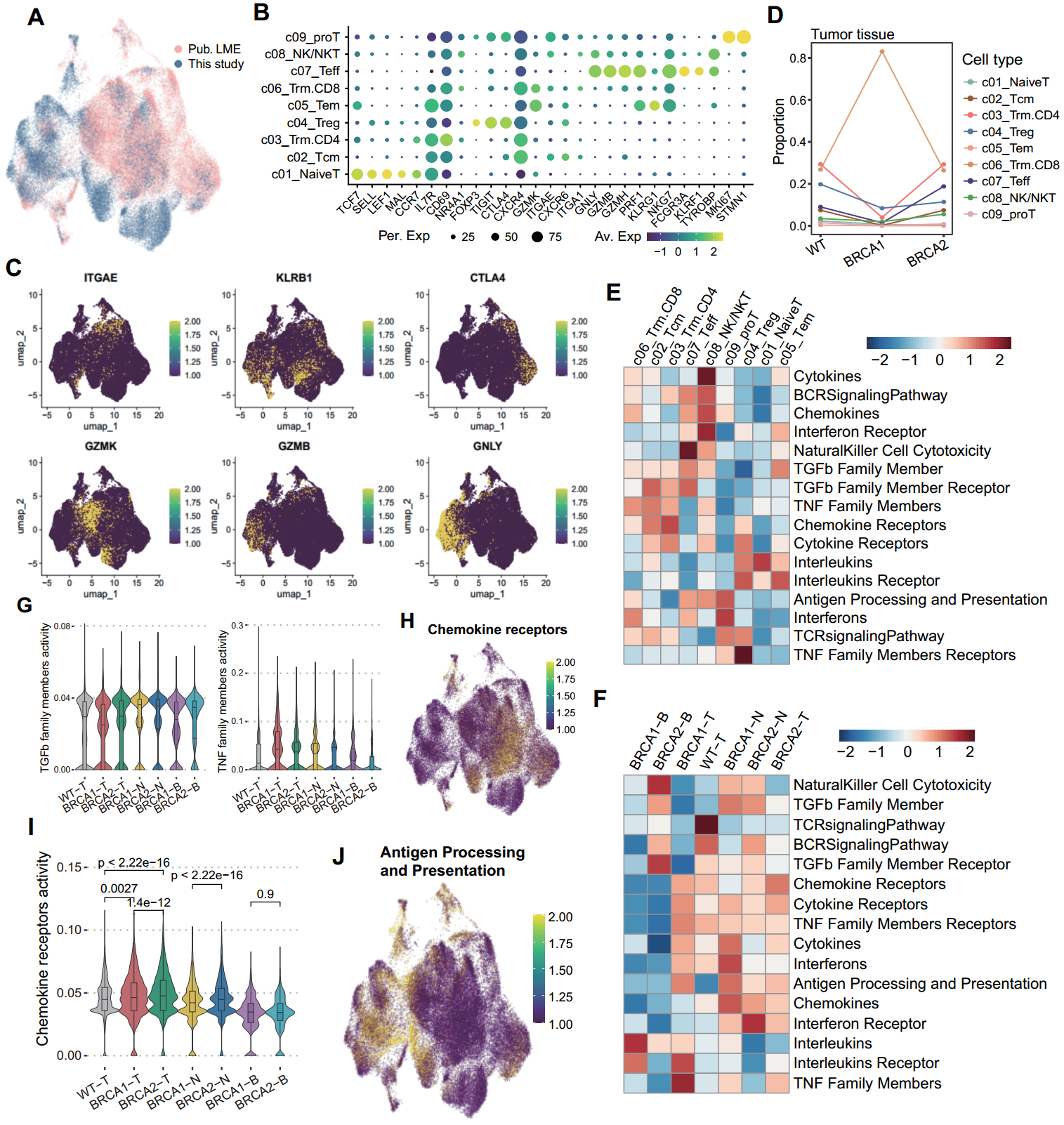

**Fig. S7. T lymphocyte activity analysis. A** UMAP visualization of data provenance in combined dataset. **B-C** The expression of the markers used for cell type identification. **D** The cell proportions of T lymphocyte subsets in wild type (WT) and *BRCA1/2* mutation groups. **E-F** Clustering analysis of immune pathway activities based on T subsets (**E**) and sample types (**F**). **G-J** Box plot (**G, I**) and UMAP plot (**H, J**) of representative immune pathways from ImmPort database.

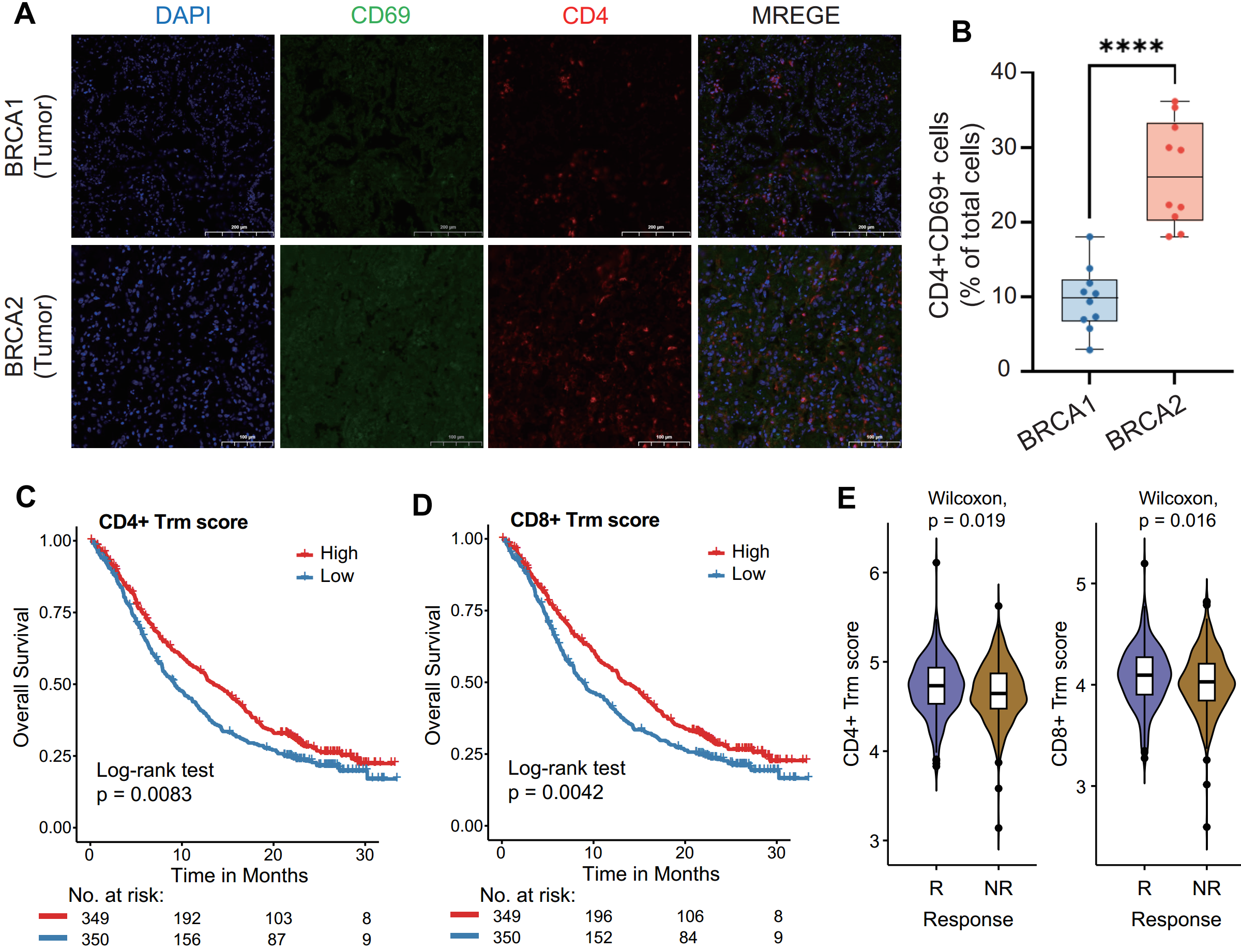

**Fig. S8. Analysis and validation of tissue-resident memory T cells. A** Multiplex immunofluorescence staining for CD4 (red), CD69 (green), and DAPI (blue) on lung tumor sections from patients with *BRCA1* and *BRCA2* mutations. **B** Quantification the relative intensity of the CD4^+^ CD69^+^ cells. The P value was computed with the two-sided Welch’s t test (ns, P>0.05; *, P<0.05; **, P<0.01; ***, P<0.001; ****, P<0.0001). **C-E** Survival analysis and response comparison (**E**) based on CD4+ Trm and CD8+ Trm score.

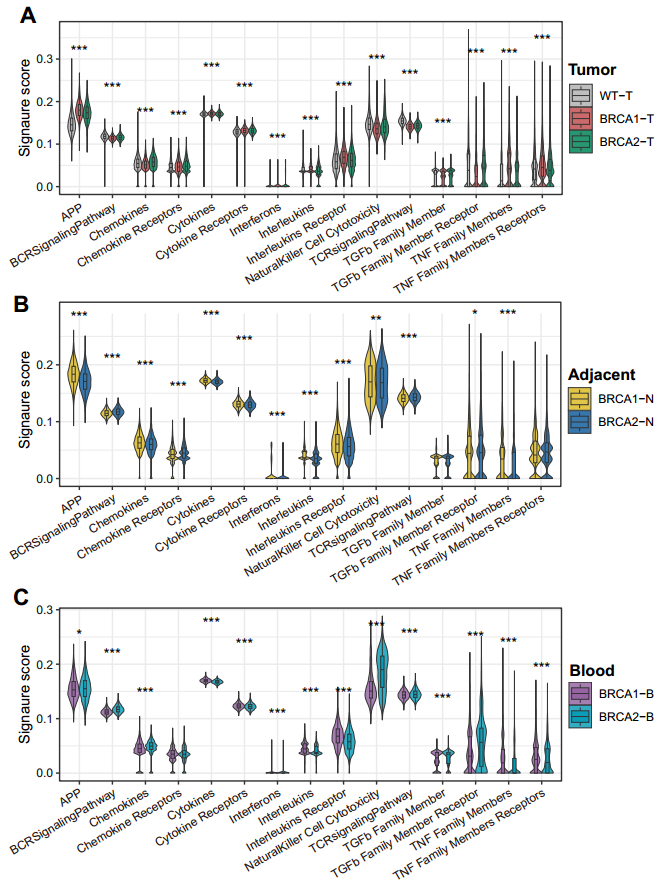

**Fig. S9. The distribution of signature score in *BRCA1/2* mutations. A-C** Comparison of immune pathway activity among wild type (WT; if any), *BRCA1* and *BRCA2* mutations in tumor (**A**), paracancerous (**B**), and blood (**C**) samples.
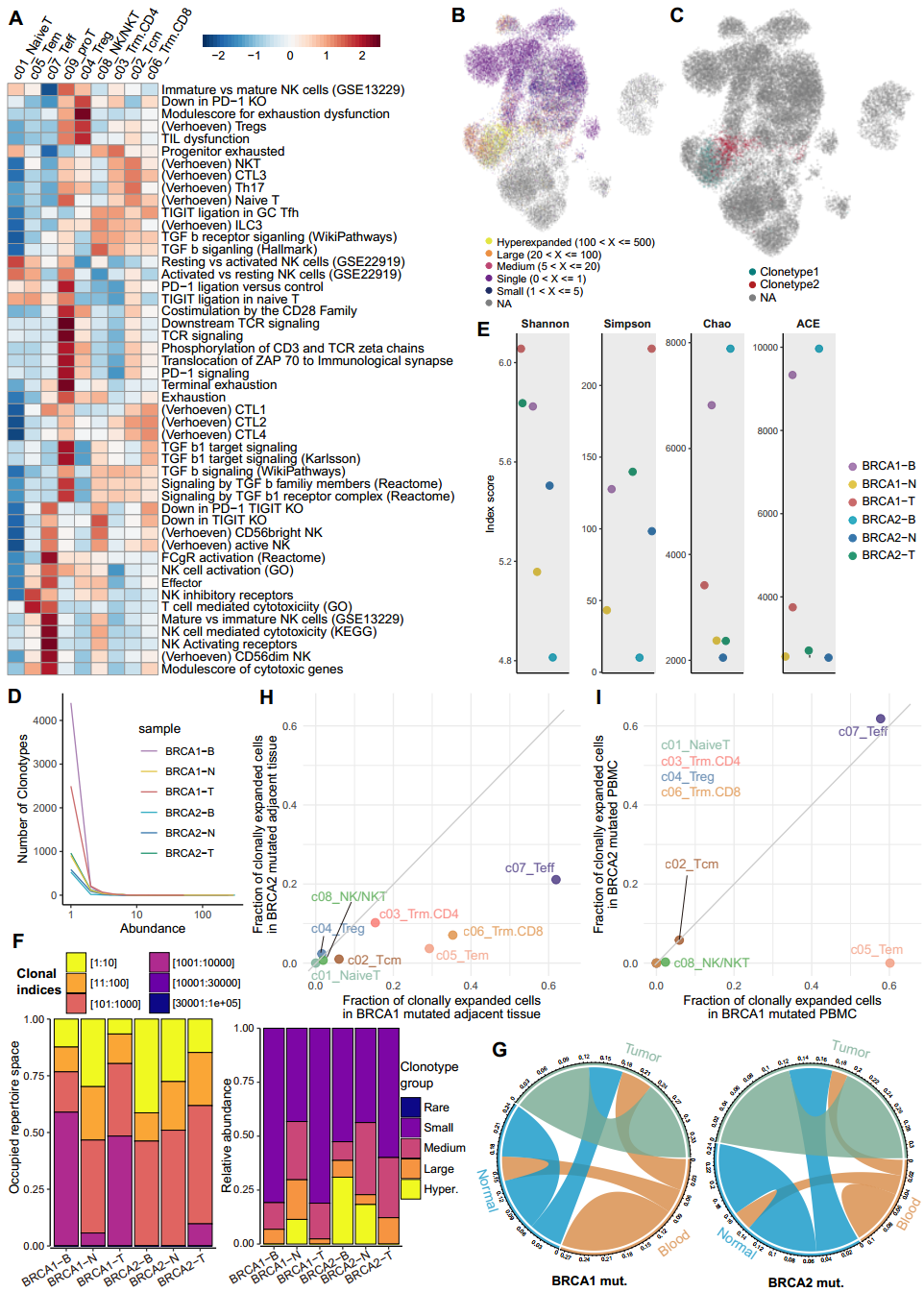

**Fig. S10. Analysis of TCR clonal expansion in T lymphocytes. A** Clustering analysis of tissue-specific T cell signature activity in T lymphocyte subsets. **B-C** UMAP visualization of expanded clonotypes (**B**) and specific clonotypes (**C**). **D** The line graph displays the number of clonotypes at specific frequencies by sample. **E** TCR diversity and TCR richness. **F** Left: The clonotypes are ranked by their frequency of occurrence, with 1:10 representing the top 10 clonotypes in each sample. Right: The proportion of different clonotypes in each sample. **G** Chord diagram visualizes the frequency of cells sharing TCRs between different samples. **H-I** The proportion of clonally expanded cells (≥5 clone size) in *BRCA1/2* mutated paracancerous (**H**) and blood (**I**) samples for specific cell types.

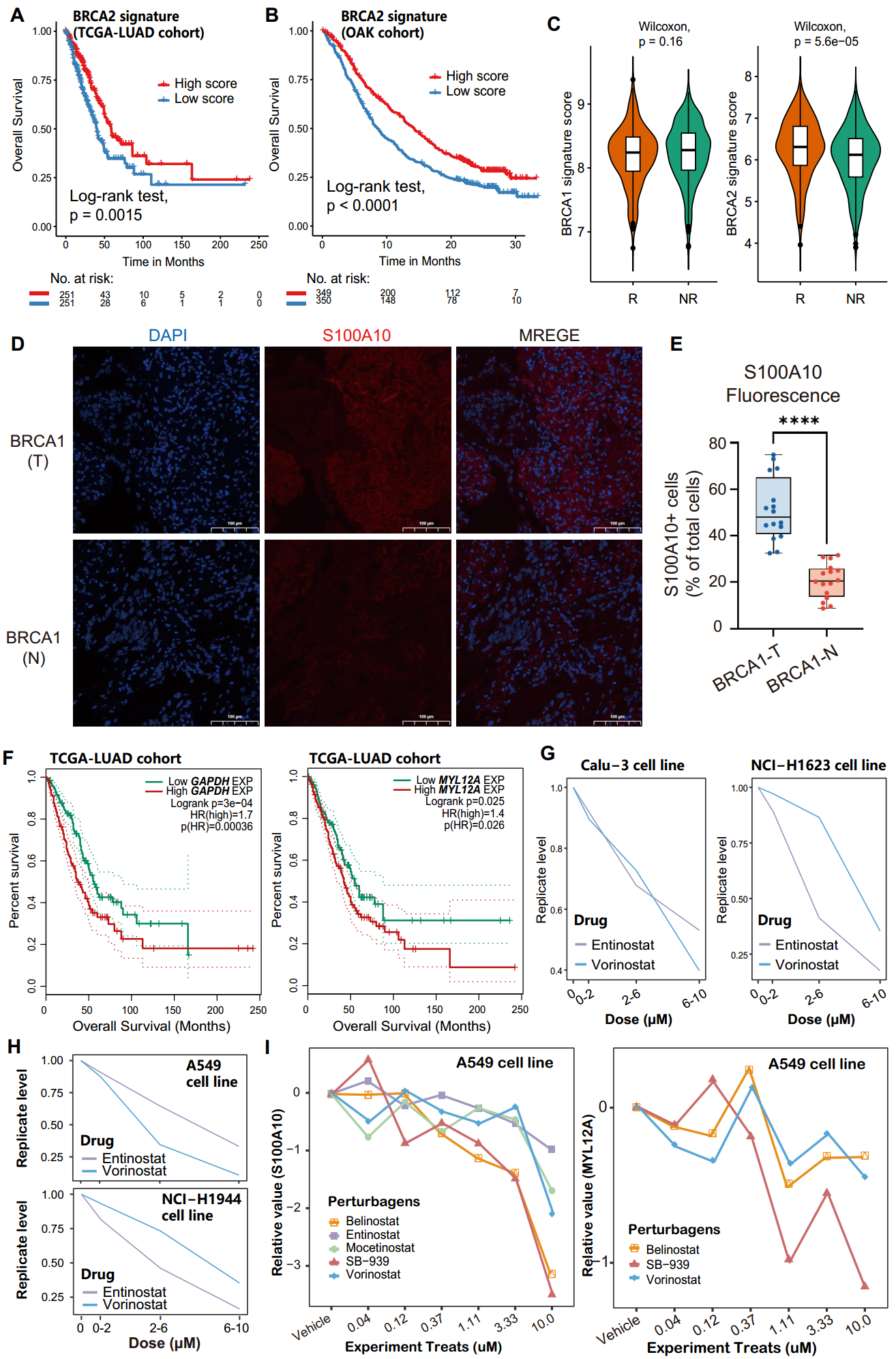

**Fig. S11. Characteristic analysis of *BRCA1/2* mutation-related signatures.** **A-B** Survival analysis of *BRCA2* mutation-related signature score in TCGA-LUAD (**A**) and OAK (B) cohorts. **C** Comparison of *BRCA1*/*2* mutation-related signature score between responders (R) and non-responders (NR) after ICB treatment. **D** Immunofluorescence staining for S100A10 (red) and DAPI (blue) on lung tumor (T) and adjacent (N) tissues from a patient with *BRCA1* mutations. Scale bars, 100 µm. **E** Quantification and estimation the relative intensity of the S100A10^+^ cells. The P value was computed with the two-sided Welch’s t test (ns, P>0.05; *, P<0.05; **, P<0.01; ***, P<0.001; ****, P<0.0001). **F** Survival analysis of *GAPDH* (left) and *MYL12A* (right) in TCGA-LUAD cohort according to the expression median. **G-H** Average cell replicate level after being treated with entinostat and vorinostat in different LUAD cell lines. **I** Expression dynamics of *S100A10* (left) and *MYL12A* (right) after treatment with different concentrations of HDAC inhibitors.

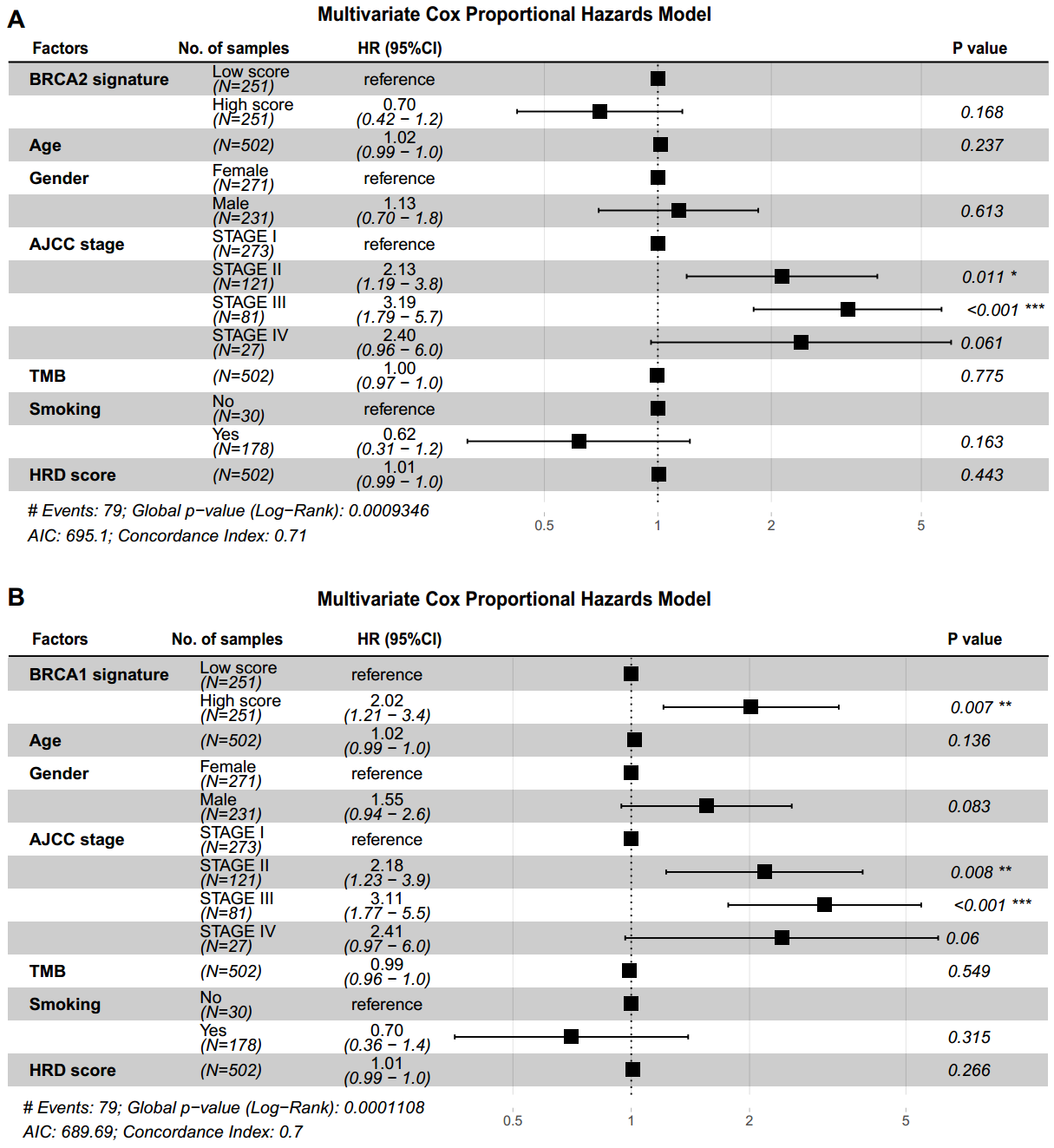

**Fig. S12. Multivariable Cox regression of the *BRCA1/2* signature score. A-B.** Multivariable Cox regression of the *BRCA2* (**A**) and *BRCA1* (**B**) signature score adjusted for age, gender, tumor stage, TMB, HRD score, and smoking status (TCGA-LUAD cohort).
