## supplementary Figure 13 for "Molecular architecture of the tumor microenvironment caused by *BRCA1* and *BRCA2* somatic mutations in lung adenocarcinoma"

**Supplementary Figure 13. Uncropped immunoblot images.**

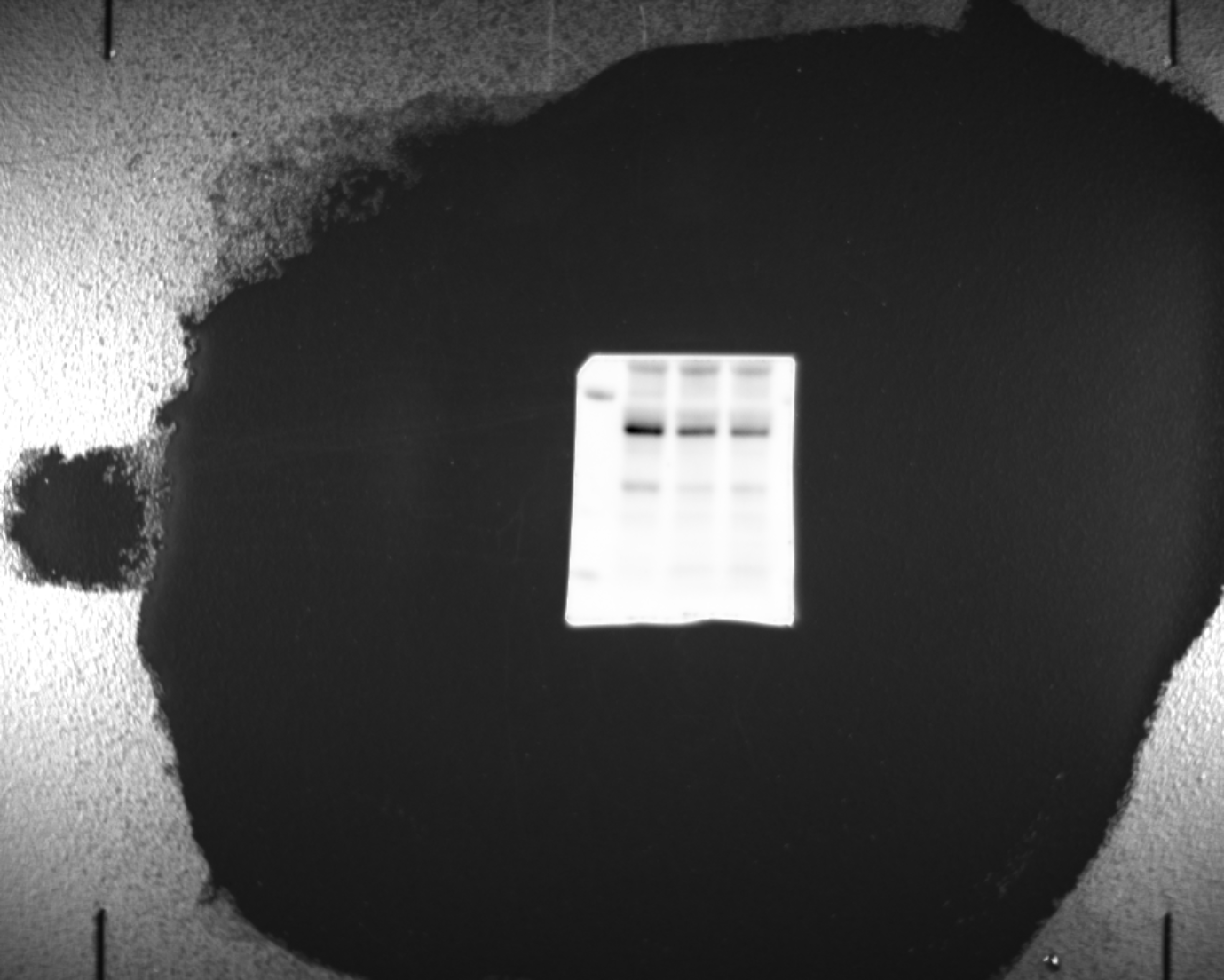
**A.** (LDHA-S100A10-GAPDH) shRNA (Corresponding to Figure 7E)

LDHA

GAPDH

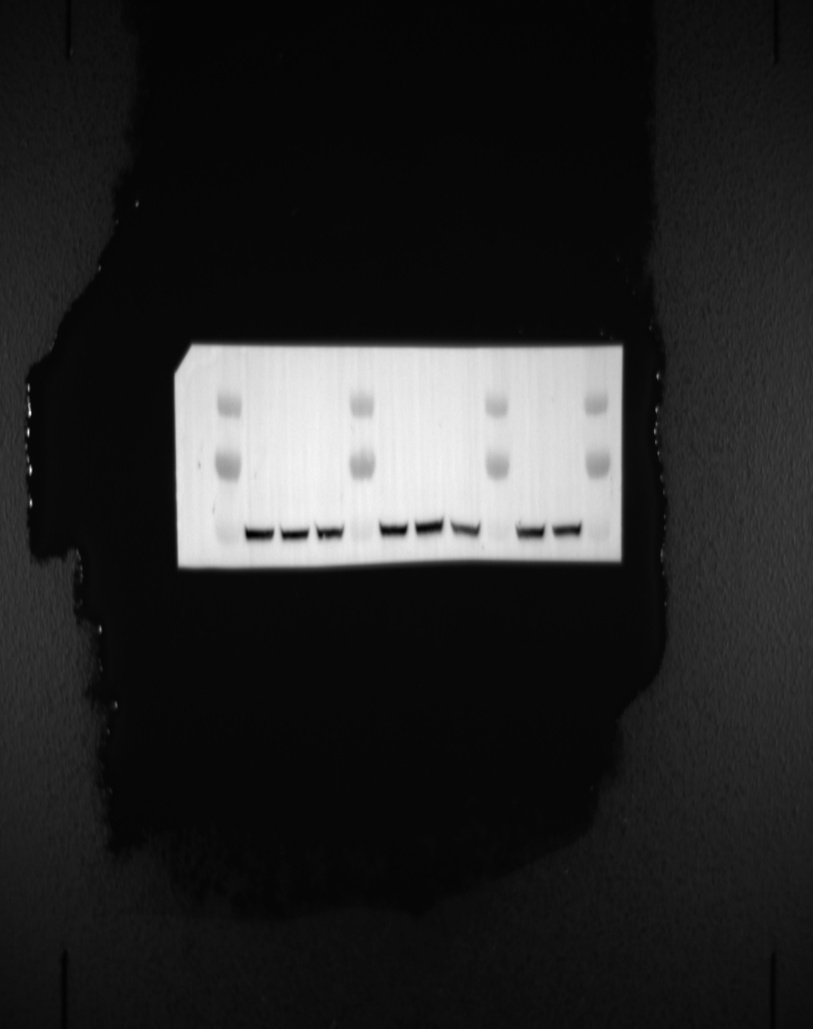

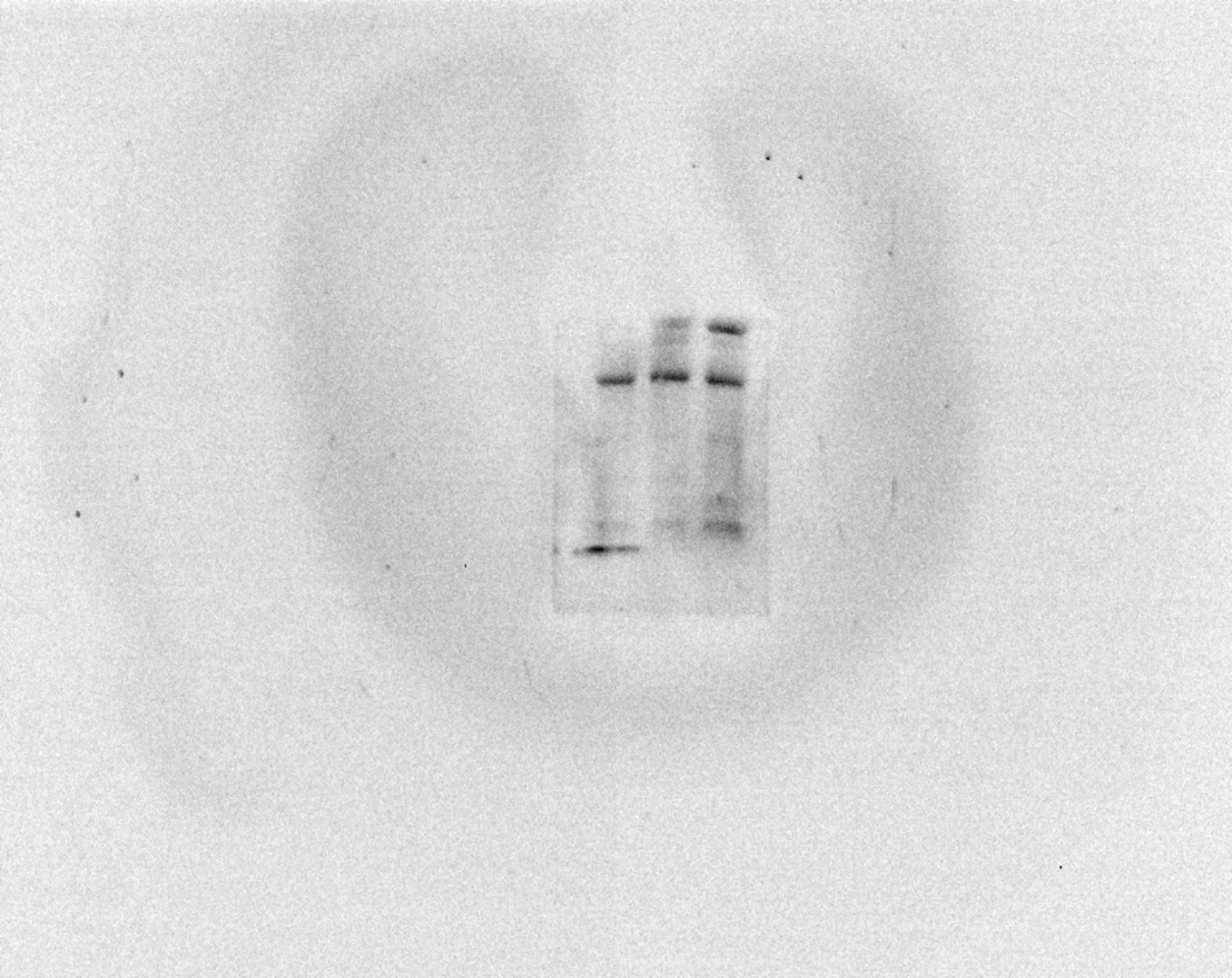

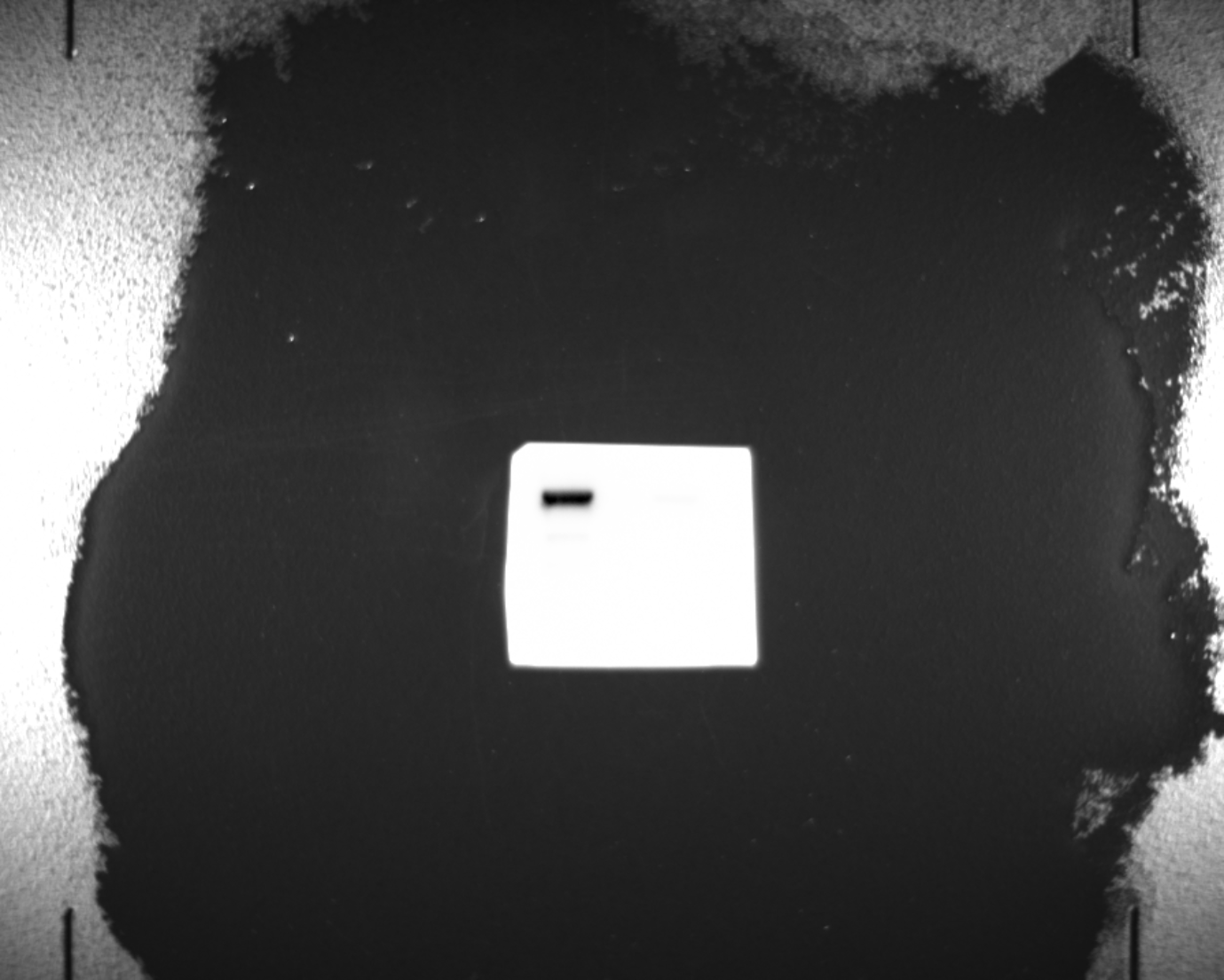

β-tubulin

S100A10

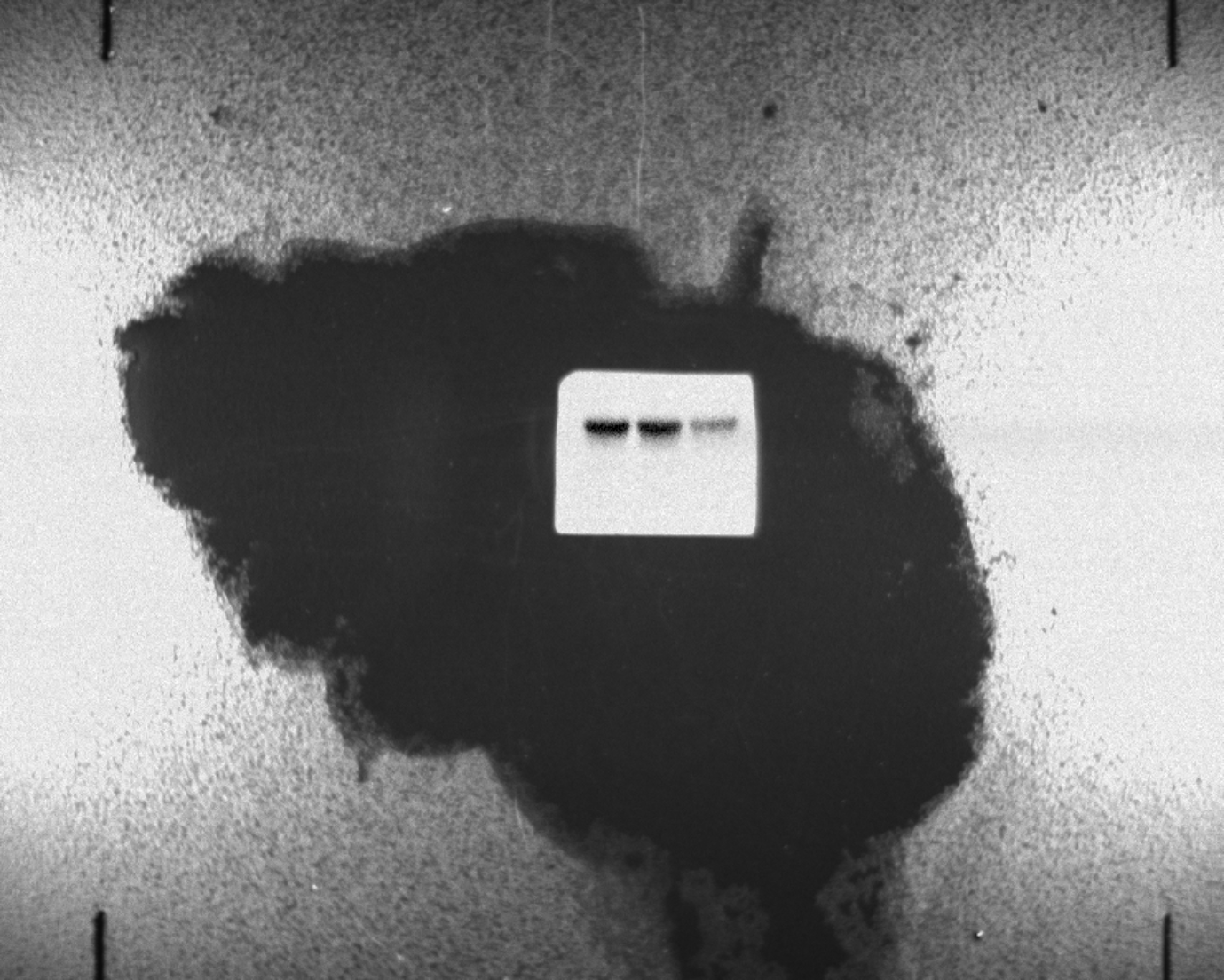
**B.** Vorinostat & Belinostat Drug treatment 48h (Corresponding to Figure 7I)

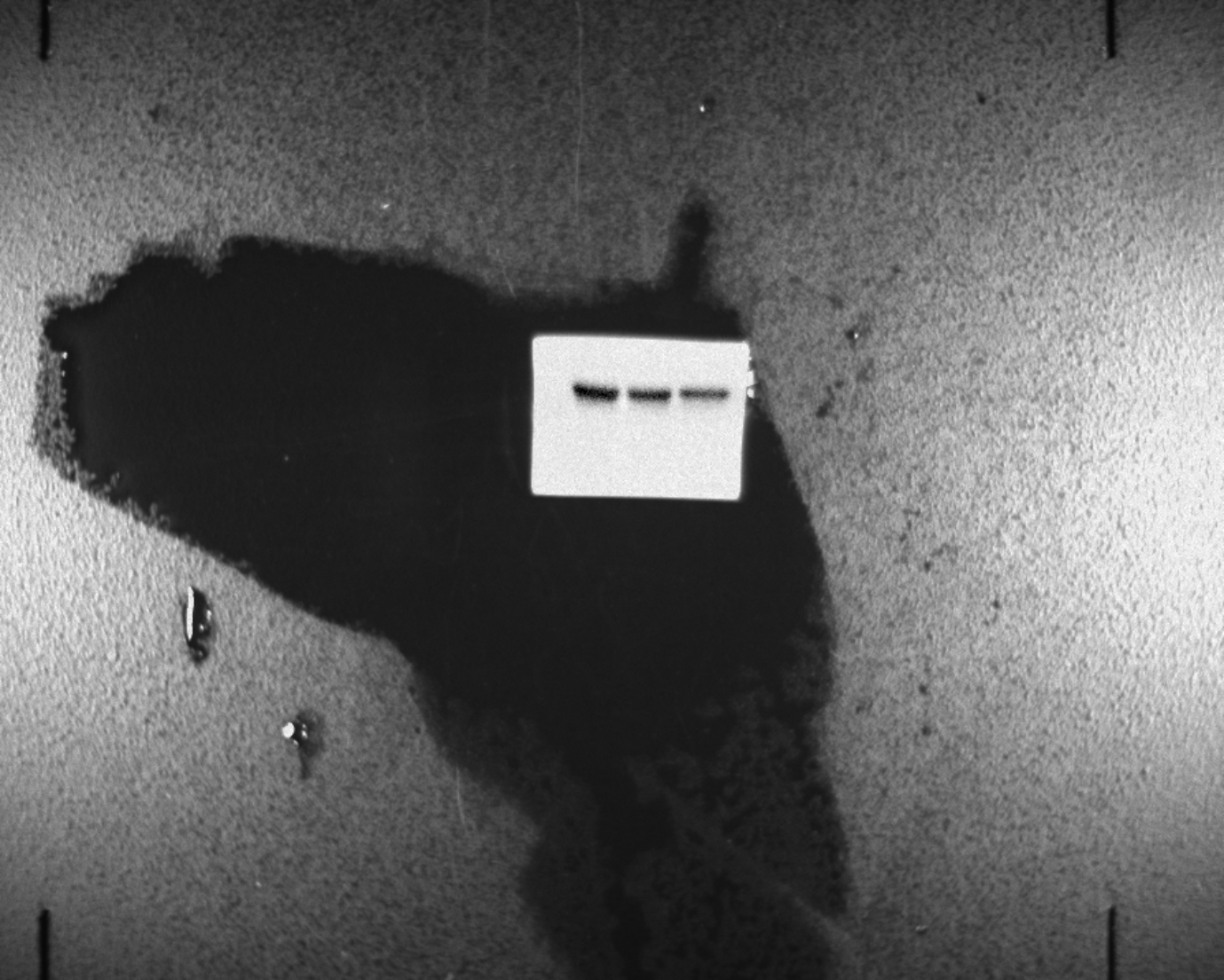

GAPDH(Belinostat)

GAPDH(Vorinostat)

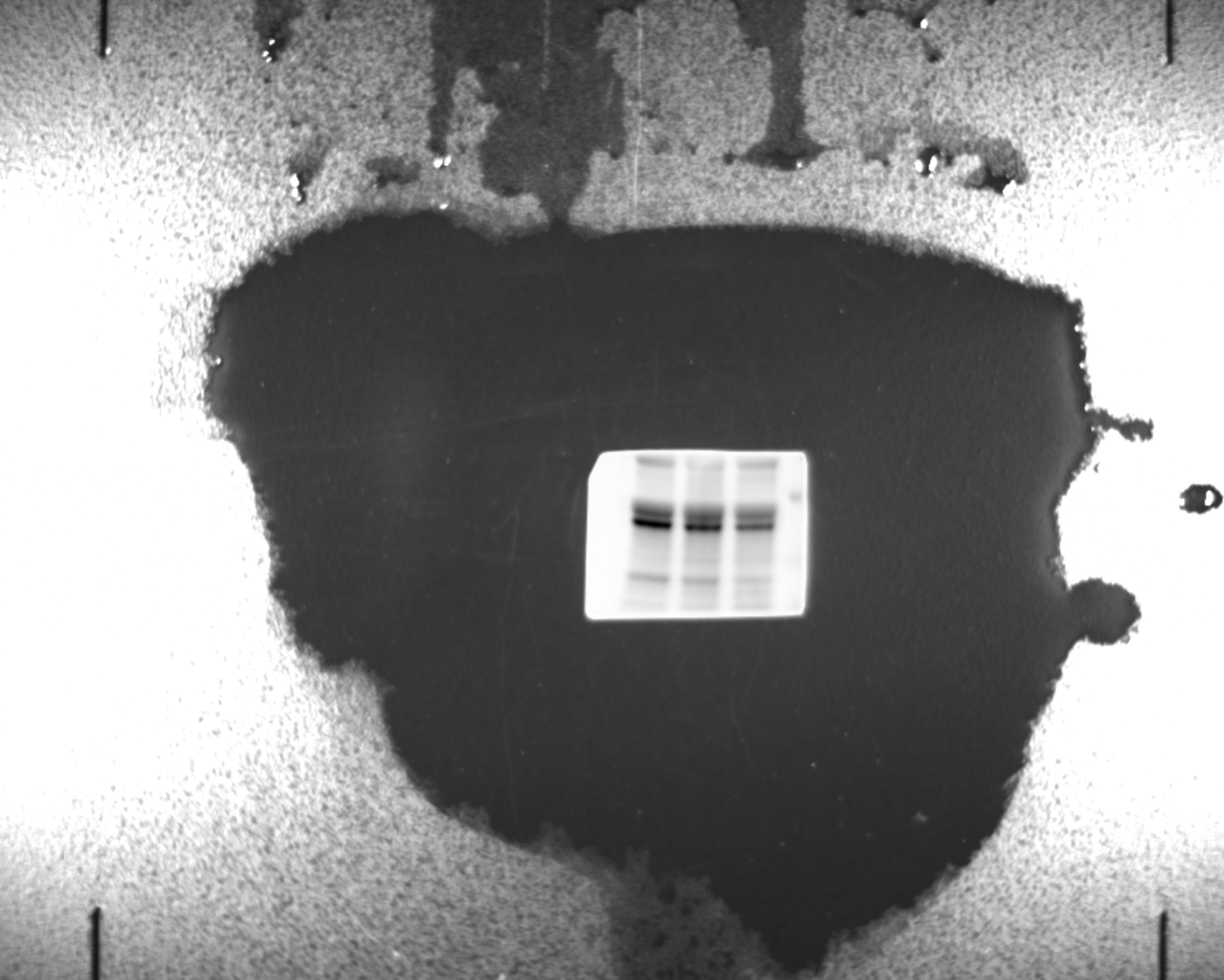

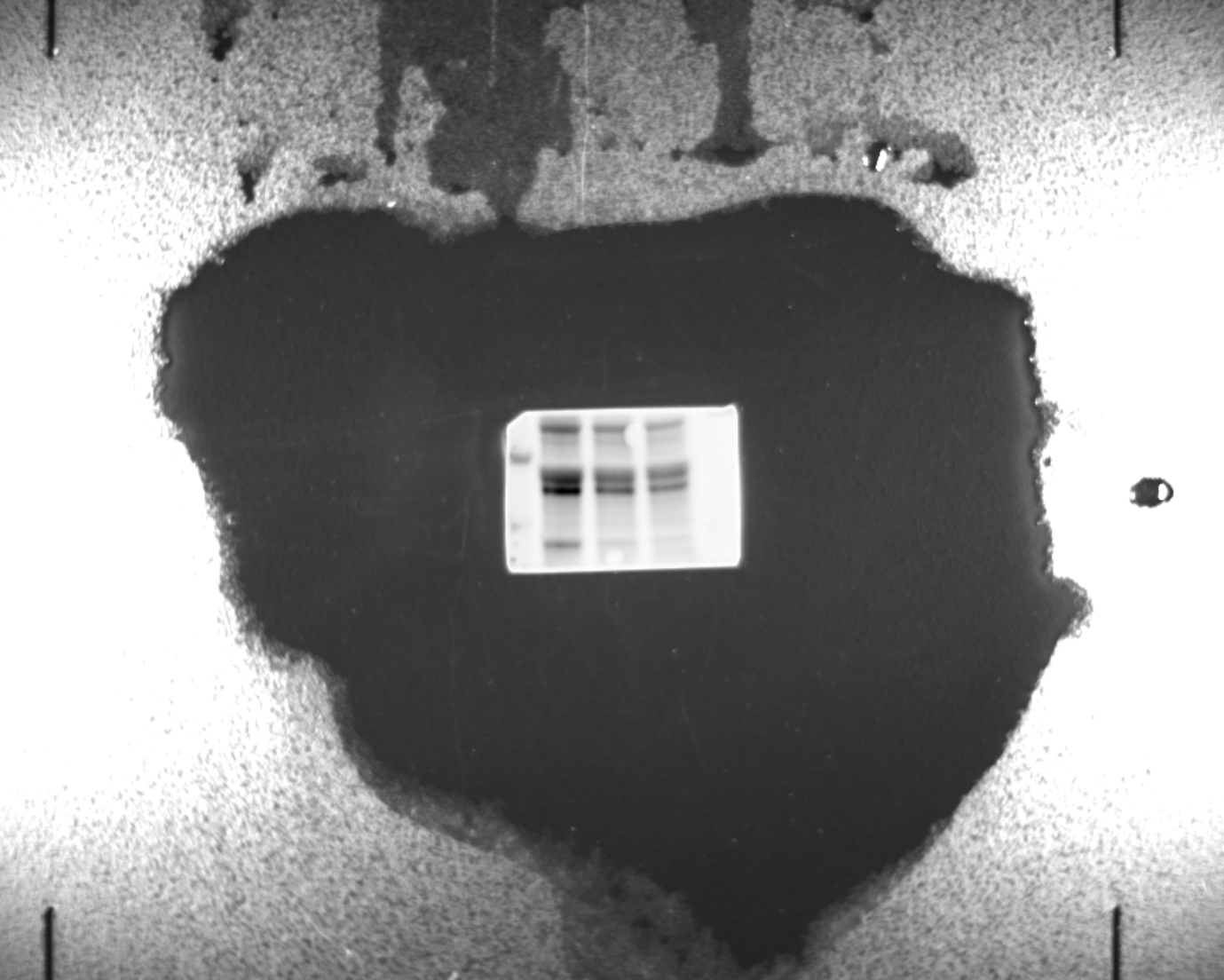

LDHA(Vorinostat)

LDHA(Belinostat)

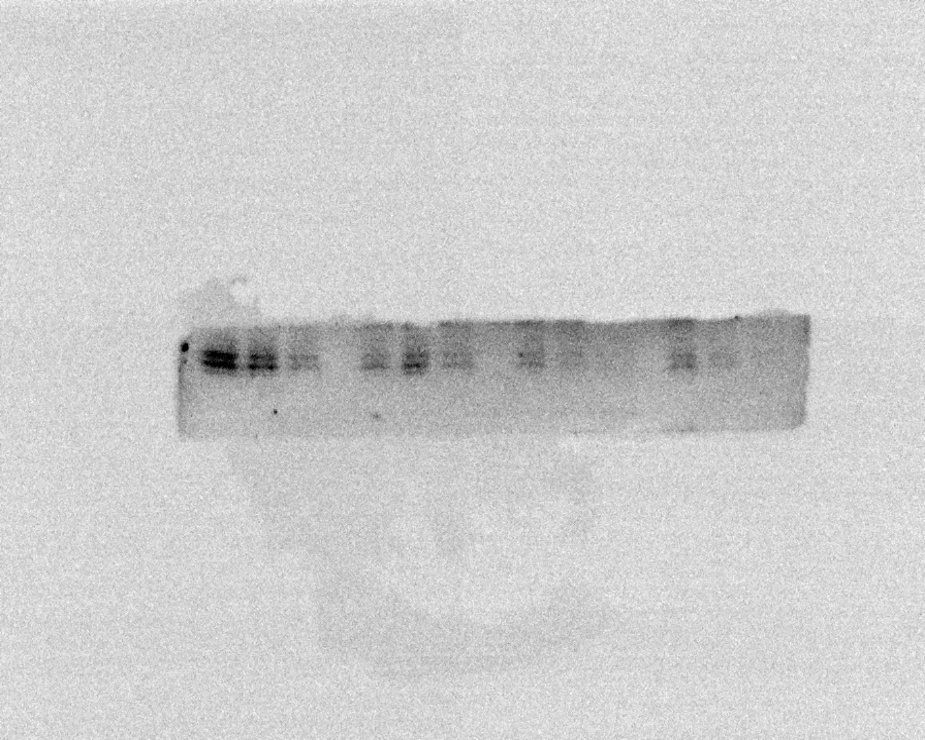

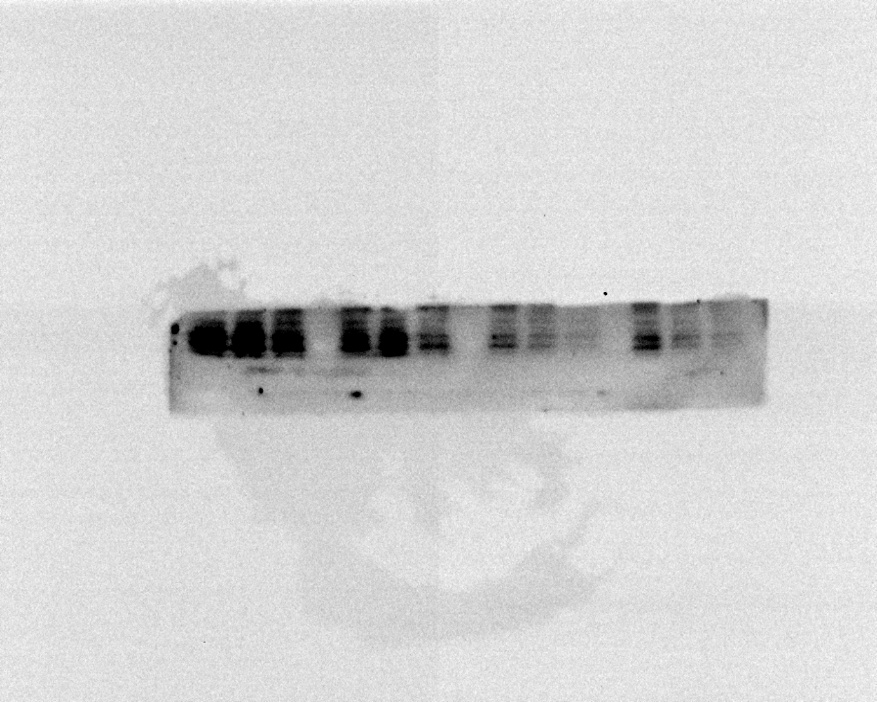

S100A10(Belinostat)

S100A10(Vorinostat)

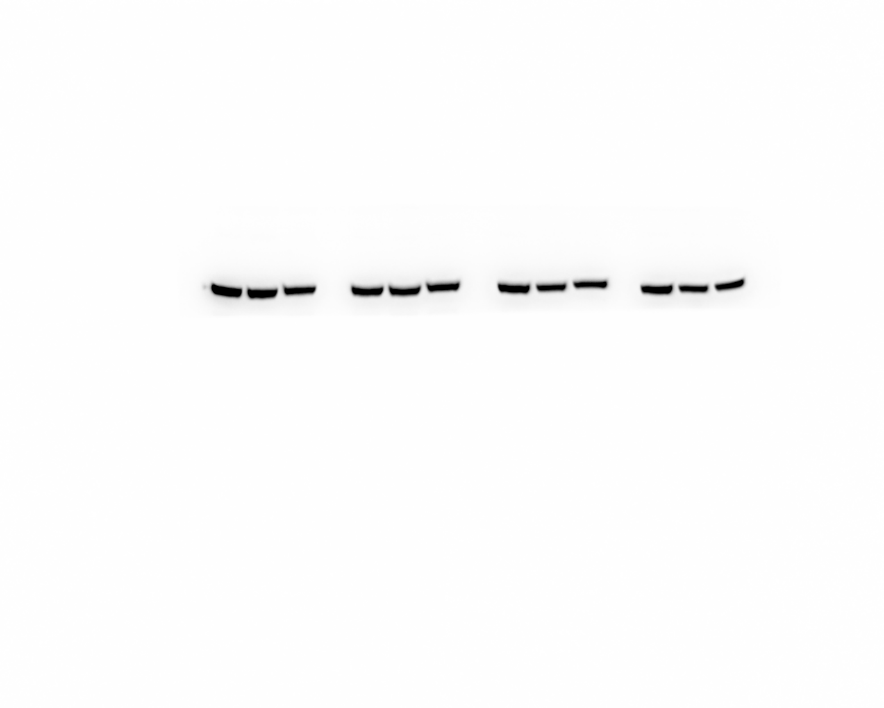

β-tubulin
